# BOSS: Beta-mixture Unsupervised Oligodendrocytes Segmentation System

**DOI:** 10.1101/2022.06.17.495689

**Authors:** Eunchan Bae, Virgilio Gonzenbach, Jennifer L Orthmann-Murphy, Russell T. Shinohara

## Abstract

To develop reparative therapies for multiple sclerosis (MS), we need to better understand the physiology of loss and replacement of oligodendrocytes, the cells that make myelin and the target of damage in MS. *In vivo* two-photon fluorescence microscopy allows direct visualization of oligodendrocytes in transgenic mouse models, and promises a deeper understanding of the longitudinal dynamics of replacing oligodendrocytes after damage. However, the task of tracking oligodendrocytes requires extensive human effort and is especially challenging in three-dimensional images. While several models exist for automatically annotating cells in two-dimensional images, few models exist to annotate cells in three-dimensional images and even fewer are designed for tracking cells in longitudinal imaging. Furthermore, the complexity of processes and myelin formed by individual oligodendrocytes can result in the failure of algorithms that are specifically designed for tracking cell bodies alone. Here, we propose a novel beta-mixture unsupervised oligodendrocyte segmentation system (BOSS) that can segment and track oligodendrocytes in three-dimensional images over time that requires minimal human input. We evaluated the performance of the BOSS model on a set of eight images obtained longitudinally. We showed that the BOSS model can segment and track oligodendrocytes similarly to a blinded human observer. Our method offers many advantages, as it does not require fully curated data, reduces computational time, and most importantly recapitulates cell dynamic patterns observed by manually tracking oligodendrocytes. Although BOSS was developed to apply to our studies on oligodendrocytes, we anticipate that it can be modified to study four-dimensional in vivo data of any brain cell.

## 1. Introduction

Oligodendrocytes are myelinating cells of the central nervous system (CNS). Myelination of axons leads to faster, more efficient communication with other neurons, and myelin loss contributes to debilitating functional deficits in multiple sclerosis (MS). The study of oligodendrogenesis and myelination processes has traditionally focused on white matter, due to the high density and alignment of myelin, allowing for *in vivo* longitudinal tracking of lesions on magnetic resonance imaging (MRI)^1^. Unlike white matter, axons are variably myelinated in the cerebral cortex^23^. Recently, a novel *in vivo* two-photon fluorescence microscopy has been developed to visualize the oligodendrogenesis and myelin sheath dynamics in cortical circuits at high resolution^4,5^. This technology allows for the investigation of the loss and replacement of oligodendrocytes and myelination patterns in acquired demyelination models^6,7^.

With development of this new imaging method, there is a growing need for automated quantification and analysis of changes in oligodendrocytes. The segmentation of oligodendrocytes involves extracting the locations of cell bodies in three-dimensional images as well as tracking oligodendrocytes over time. Currently, imaging scientists visually assess images and label the cell bodies. However, manual segmentation is costly, time-consuming and prone to inter- and intra-observer variability. Alternatively, automated segmentation would provide an efficient, consistent, scalable and reproducible way to segment the cell bodies of oligodendrocytes. Here, we propose a novel beta-mixture unsupervised oligodendrocyte segmentation system (BOSS) that can segment and track the oligodendrocytes in longitudinal three-dimensional images.

Many tools have been developed for cell segmentation in images with recent advances in machine learning. Cell-Profiler is widely used in biology, and can annotate the cell body in three-dimensional image and can track cell body in two-dimensional images, but not on three-dimensional images^8^. CellReg is a model to track neurons in calcium imaging^9^ and this model can track cell bodies in two-dimensional images, but not on three-dimensional images. Oligo-Track is a convolutional neural network (CNN) based model to track oligodendrocytes in longitudinal three-dimensional images^10^. However, this model requires a modest number of training data with full annotation. Also, this model requires extensive computing power to run (for example, an RTX 2080 Ti GPU) and takes 45 - 55 minutes per image set. Furthermore, none of the previously published methods are designed to specifically address variation in the noise levels in images (for example, across cortical depths). To address the limitations of existing tools, we developed an automated, unsupervised, fast, and human-level accurate model that needs minimal human input.

The novelty of the BOSS model is threefold. First, we modeled the intensity value of the image with mixtures of two beta distributions. The beta distribution, unlike the normal distribution, is not symmetric around the mean and thus is sufficiently flexible to model the asymmetric intensity values observed in two-photon imaging. We fitted a mixture of two beta distributions to each two-dimensional images and binarized the microscopy image into background and region of interest (ROI). For the model fitting step, we adopted an expectation-maximization (EM) algorithm^11^ that is fully unsupervised and automated. Second, we de-noised the images using a median filter while accounting for the morphology of oligodendrocye and myelin sheath. Myelin sheaths appear as weak and anisotropic noise in two-dimensional two-photon microscopy imaging. The median filter is a non-linear filter that is shown to be effective in attenuating anisotropic noise^12,13^. In our analysis, we adopted a three-dimensional window since the morphology of oligodendrocytes is spherical in shape while the myelin sheath is linear. Third, we implemented a connecting component labeling (CCL) algorithm to segment and track oligodendrocytes. CCL assigns a unique labels to all voxels in a binary image based on spatial connectedness^14^. This unsupervised labeling method has been widely used in image analysis, pattern recognition and computer vision. After aligning the three-dimensional images at each time-point by FIJI plugin ‘Correct 3D Drift’^15^, we segmented grouped voxels with same value as potential oligodendrocytes. Without a training step, which requires significant manual input, the CCL algorithm provided an automated, consistent and accurate segmentation result.

The BOSS model was developed in the open source R software environment (version 4.0.2,^16^) and the software is available in GitHub (https://pennsive.github.io/bossr/). An (a) illustration of the longitudinal course of loss (demyelination) and replacement (remyelination) of cortical oligodendrocytes and (b, c) schematic understanding the BOSS model are described in Figure 1.

**Figure 1.**
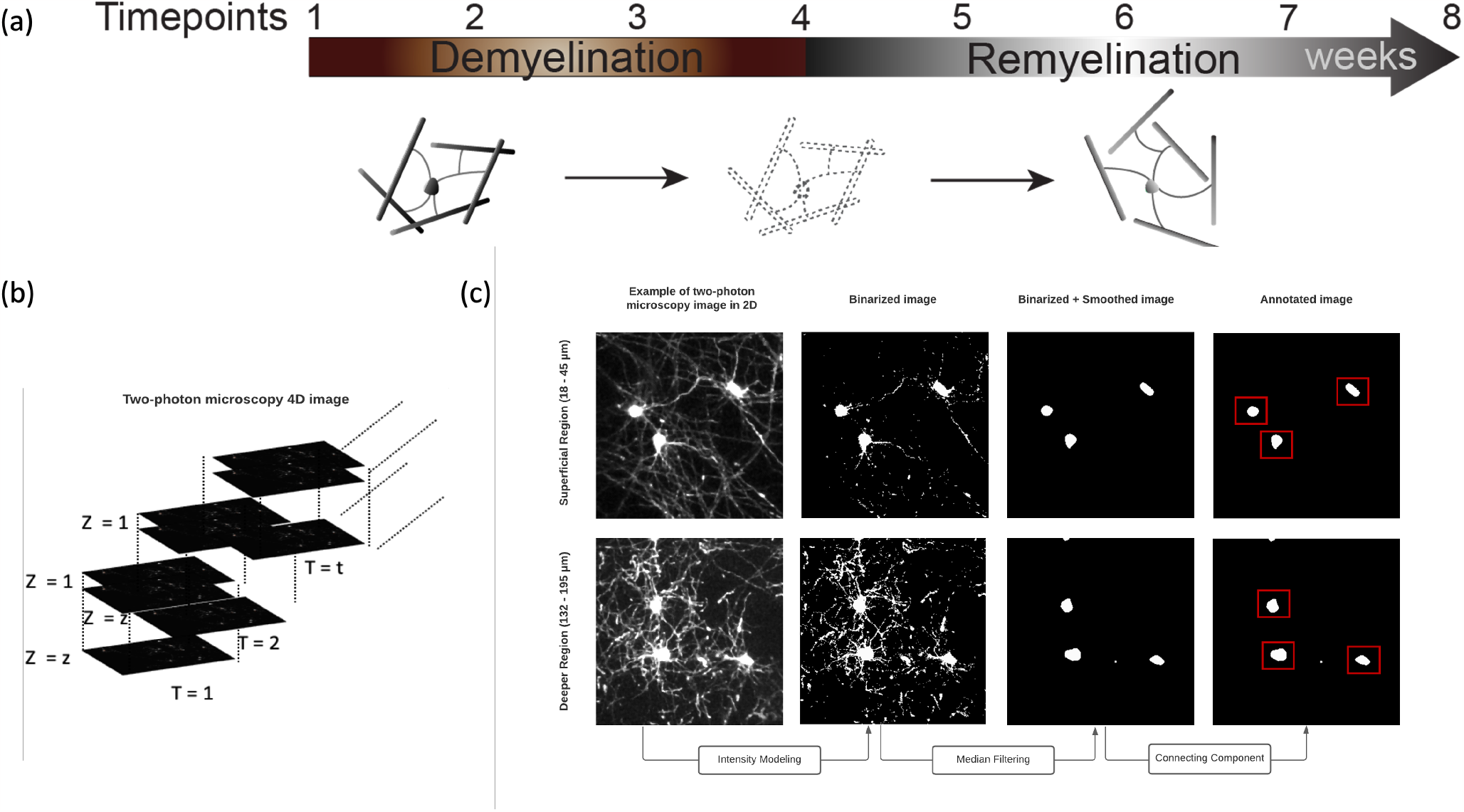
(a) An illustration of the longitudinal course of loss (demyelination) and replacement (remyelination) of cortical oligodendrocytes as previously shown (Orthmann et al 2020^6^) (b) An illustration of the raw, *in vivo* two-photon fluorescence microscopy four-dimensional images. (c) A schematic demonstrating the BOSS model for a superficial (18 - 45 µm) and deeper (132 - 195 µm) cortical region acquired by two-photon microscopy (for illustration purposes, a maximum intensity projection of each region of interest is shown). From longitudinally aligned four-dimensional images, intensity values from each two-dimensional image are modeled as a mixture of two beta distributions. From the binarized image, a median filter is applied to denoise the image. Then, connecting component labeling (CCL) is applied to annotate the oligodendrocytes. The oligodendrocytes within the red rectangle represent the BOSS annotated oligodendrocytes.

## 2. Results

We applied the BOSS model to longitudinal, *in vivo* two photon fluorescence microscopy images to visualize oligodendrocytes and astrocytes.

For visualization of oligodendrocytes, transgenic mice that express enhanced Green Fluorescent Protein (EGFP) under control of the promoter for myelin-associated oligodendrocyte basic protein (Mobp) mice were used for imaging. To determine the fate of individual oligodendrocytes in healthy versus demyelinated cortex, young adult Mobp-EGFP mice (age 8–12 weeks) were fed chow mixed with 0.2% cuprizone, a copper chelator that induces apoptosis of oligodendrocytes, or sham, for 3 weeks and then allowed to recover for at least 5 weeks. Cranial windows were placed over the somatosensory cortex, and the same region was monitored over the course of cuprizone-treatment and recovery. Throughout this article, five images from three mice treated with cuprizone and three images from two control mice were used. Details of imaging protocols are described here^6^. We excluded one time-point from control mice as there was insufficient signal-to-noise to consistently identify cell bodies with BOSS. The impact of including this image on the output of the BOSS model is discussed in 3.

While our method centers around segmenting oligodendrocytes, this can be extended to study other types of cell bodies. For example, astrocytes are complex cells distributed throughout the central nervous system (CNS), with highly branched processes that interact with all other CNS cells^17^. Astrocytes have many important functions in development and the healthy adult CNS, but also respond and contribute to CNS damage^18^. To demonstrate the applicability of the BOSS model to astrocytes, we applied the BOSS model to a set of similarly acquired four-dimensional images from transgenic mice expressing TdTomato in the cytosplasm of astrocyte and EGFP in oligodendrocytes (GLAST-CreER; R26-lsl-tdTomato; MOBP-egfp) and annotated the astrocyte over time. We fitted the BOSS model with the parameters used for oligodendrocyte annotation as astrocytes and oligodendrocytes look similar.

To evaluate the performance of the BOSS model, we compared it with annotations conducted previously by blinded human observers as previously reported for these images^6^. For oligodendrocytes, the number of oligodendrocytes as well as the number of new oligodendrocytes and lost oligodendrocytes were annotated. To account for correlation across the multiple time points, we fitted a linear mixed effect model with the number of annotated oligodendrocytes by blinded human observes as the outcome, the number of annotations by the BOSS model as a fixed effect, and each image set as a random effect. An estimated coefficient greater than one implies the under-annotation whereas an estimated coefficient less than one implies the over-annotation. The model was fit and fixed effects as well as the profile likelihood-based confidence interval (CI) were calculated using *lmerTest::lmer()* in R^19^. For astrocytes, the number of astrocytes as well as the number of new oligodendrocytes and lost oligodendrocytes were annotated.

### 2.1 Oligodendrocytes Annotation Results of the BOSS model across time-points

We evaluated the BOSS model using annotations conducted by blinded human observers as the gold standard. The bar graphs of proportions of oligodendrocytes in a 425 µm x 425 µm x 300 µm volume superficial region of somatosensory cortex compared to the baseline (week 1), as well as the number of new/lost oligodendrocytes annotated by the BOSS model and the blinded human observer, are presented in Figure 2. The Bland-Altman (BA) plots are also presented in Figure S2 to assess the agreement between two quantitative methods of measurement^20^.

**Figure 2.**
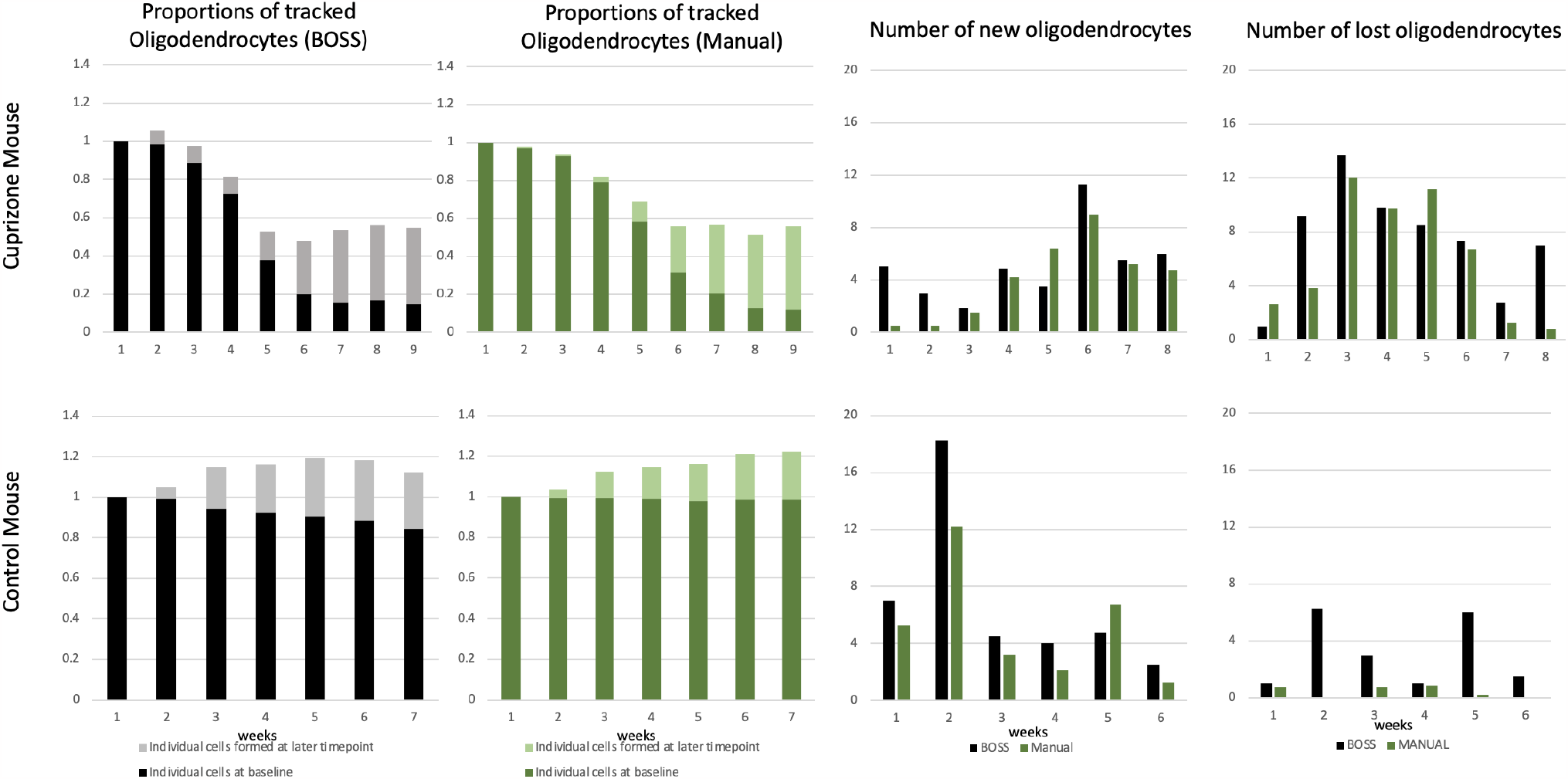
Results of the BOSS model across time-points. The first row shows the result of the BOSS model when applied in cuprizone-treated mice. The second row shows the result of the BOSS model when applied in control mice. The first/second column shows the proportions of the number of tracked oligodendrocytes by the BOSS model/blinded human observer compared to the week 1. The third column shows the average number of newly formed oligodendrocytes at each time-point. Each time-point represents the number of new oligodendrocytes observed at its next time-point. The fourth column shows the average number of lost oligodendrocytes at each time-point. Each time-point represents the number of lost oligodendrocytes observed at its next time-point.

For images from cuprizone-treated mice, the BOSS model annotated the number of oligodendrocytes similarly to the blinded human observer compared to its baseline. The estimated regression coefficient of the BOSS model on blinded human observe was 1.06 (95% CI: 0.77 to 1.28). The BOSS model over-annotated the number of new oligodendrocytes over time-points and the estimated regression coefficient of the BOSS model for the blinded human observe but the difference was minimal (mean absolute difference: 1.83). Similarly, the BOSS model over-annotated the number of lost oligodendrocytes over time-points, but the difference was minimal (mean absolute difference: 2.45). Both the BOSS model and the blinded human observer reported the maximum number of new/lost oligodendrocytes at the same time-point. Both reported the maximum number of the number of new oligodendrocytes at sixth time-point (corresponding to 1 week recovery) and lost oligodendrocytes at third time-point (2 weeks of cuprizone treatment). BA plots (in first row) show the good agreement between the BOSS model and the blinded human observer as almost all data points are within the 1.96 times the standard deviance of the differences. BA plots of the new/lost oligodendrocytes show the unbiased measurement of the BOSS model compared to the manual annotation.

For images from control mice, the BOSS model annotated the number of oligodendrocytes similarly to the blinded human observer compared to its baseline. The estimated regression coefficient of the BOSS model on blinded human observer was 1.09 (95% CI: 0.90 to 1.27). The BOSS model over-annotated the number of new oligodendrocytes over time-points, but the difference was minimal (mean absolute difference: 2.35). The BOSS model over-annotated the number of lost oligodendrocytes over time-points, but the difference was minimal (mean absolute difference 2.69). BA plots (in second row) show good agreement between the BOSS model and the blinded human observer as all data points are within the 1.96 times the standard deviance of the differences. BA plots (in second row) show good, unbiased agreement between the BOSS model and the blinded human observer as all data points are within the 1.96 times the standard deviance of the differences.

### 2.2 Oligodendrocytes Annotation Results of the BOSS model across time-points, stratified by the depth of somatosensory cortex

We next assessed the performance of the BOSS model by stratifying by depth of superficial somatosensory cortex in increments of 100 µm from the pial surface as previously analyzed in Orthmann-Murphy et al^6^. The proportions of tracked oligodendrocytes as well as the count of new/lost oligodendrocytes as annotated by the BOSS model and blinded human observer, stratified by depth, are presented in Figure 3 for cuprizone-treated mice and in Figure S3 for control mice.

**Figure 3.**
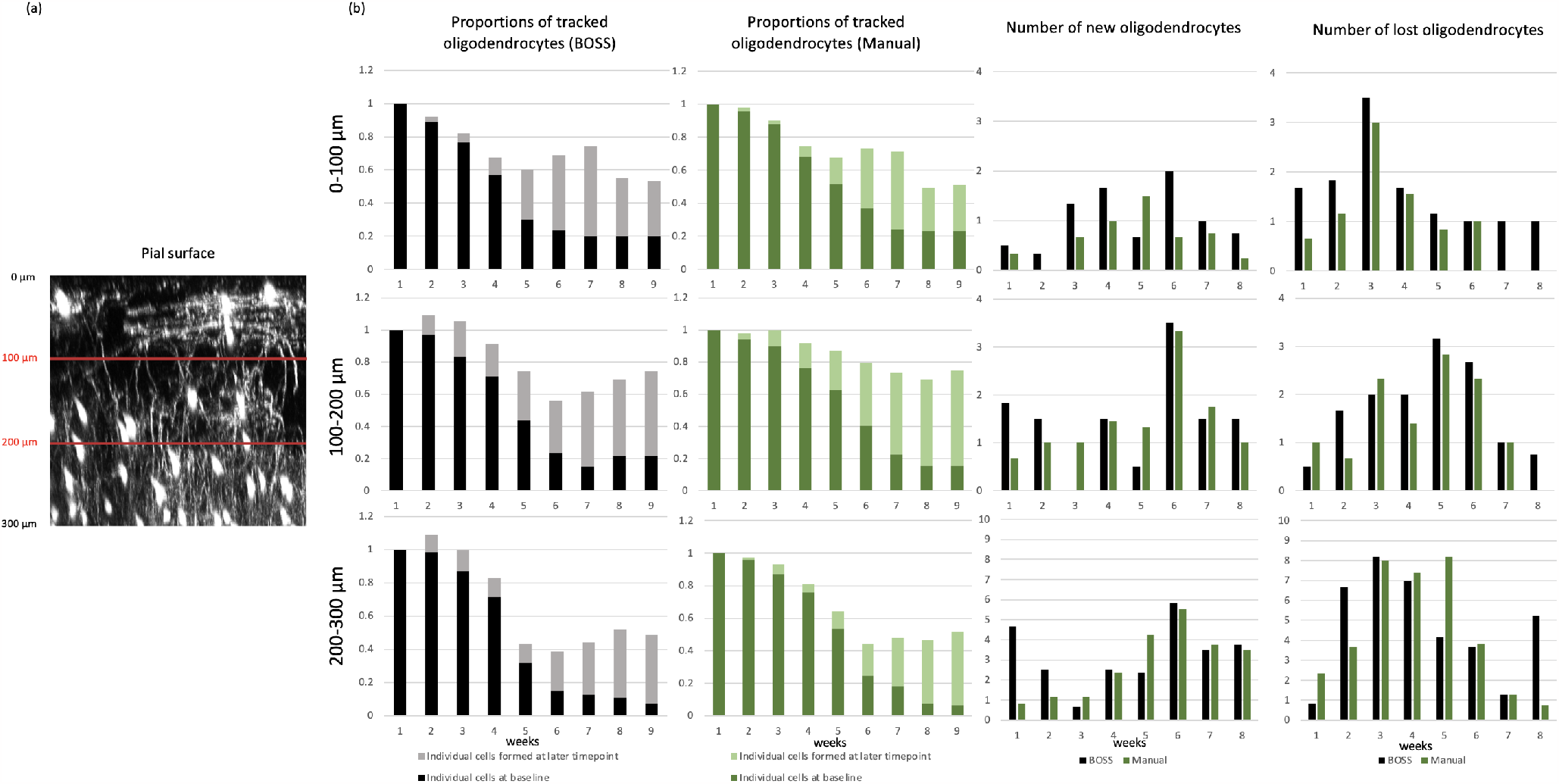
(a) An illustration of the image showing oligodendrocytes with maximum intensity Y-projection, separated by red lines at intervals of 100 µm. (b) Results of the BOSS model across time-points stratified by depth in somatosensory cortex for cuprizone-treated mice. The rows indicate the 0-100 µm, 100-200 µm, and 200-300 µm depth regions relative to pial surface. The first column shows the proportions of the number of tracked oligodendrocytes by the BOSS model compared to the week 1. The second column shows the proportions of the number of tracked oligodendrocytes by blinded human observer compared to the week 1. The third column shows the average number of newly formed oligodendrocytes. The rightmost column shows the average number of lost oligodendrocytes.

In all zones of images from cuprizone-treated mice, the annotation by the BOSS model corresponded to the blinded human observer with significant *p*-values. In the 0-100 µm zone (superficial), the estimated regression coefficient of the BOSS model on blinded human observer was 0.84 (95% CI: 0.63 to 1.04). In the 100-200 µm zone, the estimated regression coefficient of the BOSS model on blinded human observer was 0.53 (95% CI: 0.39 to 0.68). In the 200-300 µm zone, the estimated regression coefficient of the BOSS model on blinded human observer was 1.08 (95% CI: 0.98 to 1.18). The BOSS model over-annotated new/lost oligodendrocytes in 0-200 µm zones, the superficial region with relatively fewer oligodendrocytes^4^ and under-annotated new/lost oligodendrocytes in 200-300 µm zones, where density of oligodendrocytes is higher^4^, compared to the blinded human observer, but the difference was minimal. In the 0-100 µm zones, the mean absolute difference of the number of annotated new oligodendrocytes was 0.59 and the mean absolute difference of the number of annotated lost oligodendrocytes was 0.58. In the 100-200 µm zones, the mean absolute difference of the number of annotated new oligodendrocytes was 0.56 and the mean absolute difference of the number of annotated lost oligodendrocytes was 0.58. In the 200-300 µm zones, the mean absolute difference of the number of annotated new oligodendrocytes was 1.07 and the mean absolute difference of the number of annotated lost oligodendrocytes was 1.72.

For images from control mice in the 0-100 µm zones, the estimated regression coefficient of the BOSS model on blinded human observer was 1.07 (95% CI: 0.99 to 1.15). In the 100-200 µm zones, the estimated regression coefficient of the BOSS model on blinded human observer was 0.86 (95% CI: 0.69 to 1.00). In the 200-300 µm zones, the estimated regression coefficient of the BOSS model on blinded human observer was 0.97 (95% CI: 0.58 - 1.28). The BOSS model over-annotated new/lost oligodendrocytes in almost all zones compared to the blinded human observer, but the difference was minimal. In the 0-100 µm zones, the mean absolute difference of the number of annotated new oligodendrocytes was 0.60 and the mean absolute difference of the number of annotated lost oligodendrocytes was 0.21. In the 100-200 µm zones, the mean absolute difference of the number of annotated new oligodendrocytes was 0.52 and the mean absolute difference of the number of annotated lost oligodendrocytes was 0.54. In the 200-300 µm zones, the mean absolute difference of the number of annotated new oligodendrocytes was 2 and the mean absolute difference of the number of annotated lost oligodendrocytes was 2.08.

### 2.3 Astrocytes Annotation Results of the BOSS model across time-points

The result of the BOSS model is presented in Fig 4 and the raw images, images with the BOSS-annotation as well as the spreadsheet that reports the coordinates of cell bodies, unique cell identifier, and the size of the cell bodies can be found here. The result showed that the BOSS model annotated the astrocytes similarly to the blinded human observer (Mean absolute difference: 5.2). The BOSS model over-annotated the new/lost cells at a later time-point in Mouse II and we confirmed that this was due to differences in signal-to-noise ratio of acquired TdTomato signal at T=7 and T=8. As the BOSS model was developed to identify dying oligodendrocytes as exhibiting decreasing intensity and new cells with increasing intensity, the BOSS model reported loss of cells at T=7 and gain of cells at T=8, where signal-to-noise was greater. Overall, this result shows that the BOSS model can be extended to annotate the other cell types with default model parameters, but also reflects distinct biological parameters unique to the cell being tracked, as well as opportunities to modify BOSS for different cell types. For instance, we can consider using lenient parameters as astrocytes do not produce myelin sheaths. Choosing the candidate voxels with intensity values up to 98th percentile instead of 99th percentile in the pre-processing step, and setting the 70th percentile of intensity value as a threshold instead of 80th percentile to label the ROI and background when the two beta distributions are not found, improved the mean absolute difference of astrocyte annotation between the BOSS model and the blinded human observer by 0.9 (5.2 → 4.3).

**Figure 4.**
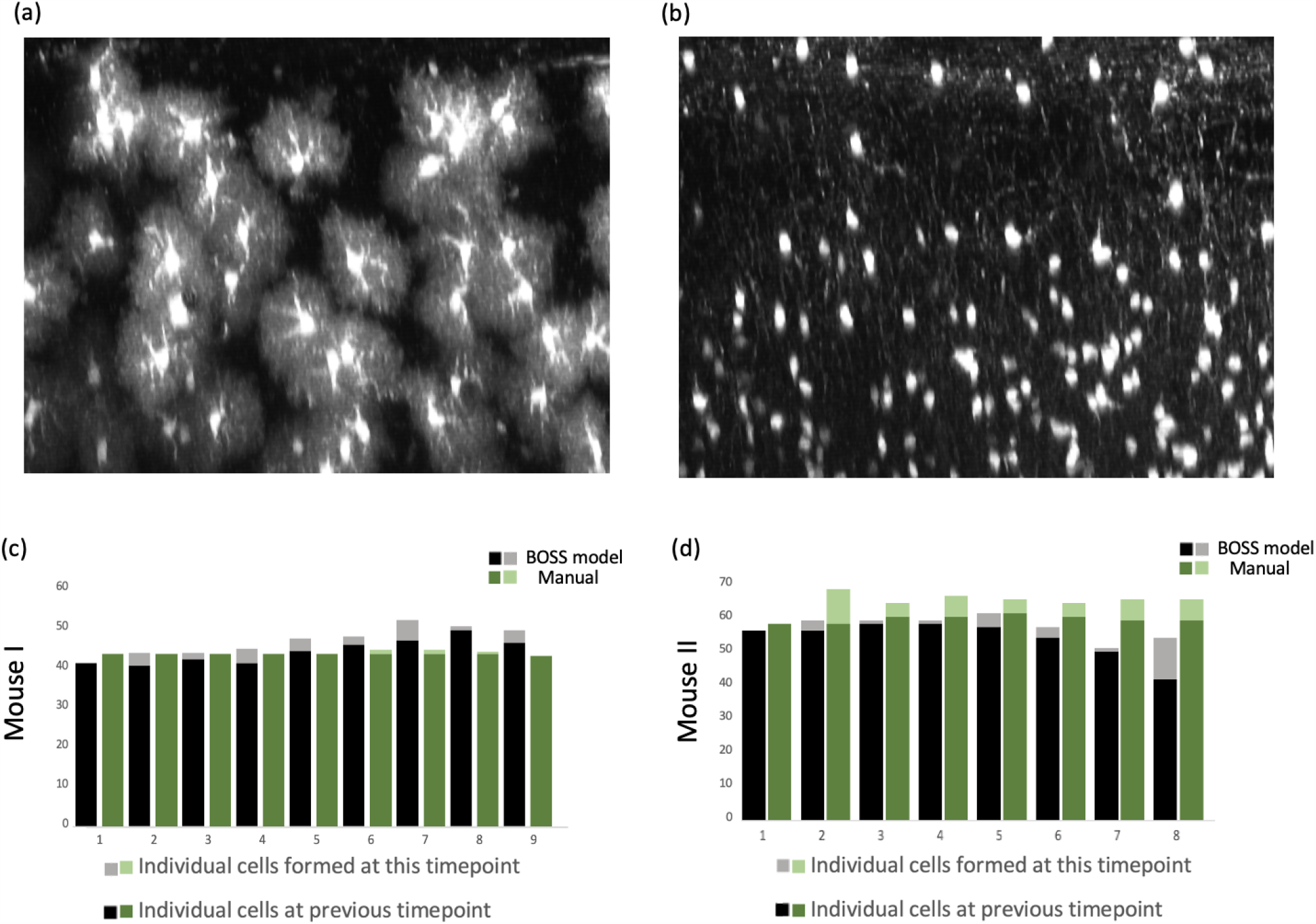
(a) An illustration of the image of astrocyte with Z-projection (top-down view, 405 µm x 300 µm) (b) An illustration of the image of astrocyte with maximum intensity Y-projection (coronal view, 405 µm x 300 µm x 425 µm). (c,d) Results of the BOSS model to astrocytes across time-points. The number of tracked astrocytes across time-points, annotated manually at the previous time-point (by the BOSS model in black, by the blinded human observer in green) and each time-point (by the BOSS model in light grey, by the blinded human observer in light green) is presented.

## 3 Discussion

We propose an unsupervised, fast, and accurate oligodendrocyte segmentation system that segments and tracks oligodendrocytes in longitudinal three-dimensional images with minimal human input. The BOSS model adjusts for the heterogeneous brightness of image across cortex depth and removes myelin sheaths and noise in order to segment the cell bodies of oligodendrocytes. By using CCL, the BOSS model segments and tracks the oligodendrocytes with volumetric quantification of each oligodendrocytes.

The BOSS model was developed so that no training step is needed. This achievement is crucial as manual cell segmentation can lead to long delays in analysis. In our study, it took approximately six hours to manually annotate and track oligodendrocytes for a single image set used in this analysis by human observer. Also, it could take much more time if a user wants to use a supervised method such as convolutional neural network (CNN) that requires a fully annotated training set as well as model tuning. As the BOSS model was developed as an unsupervised method, we expect our BOSS model can solve this bottleneck. In addition, unlike deep learning technologies which have been criticized for its black-box nature, the BOSS model allows a user to understand each step and can therefore tune parameters.

The BOSS model showed unsatisfactory performance when applied to the control mouse data set that included images with poor signal-to-noise (Fig S4). Based on manual tracking, we expected to see a stable number of oligodendrocytes with increasing the number of new oligodendrocytes^6,4,5^. However, the BOSS model reported loss of oligodendrocytes at a time-point with low signal-to-noise of the GFP signal. This was due to the variability of image quality intrinsic to longitudinal *in vivo* imaging, including varying dural thickness over weeks^21^. Supplementary figure in Figure S5 in the first row shows the one of original images that represent the varying image acquisition parameters. At T=8, the GFP signal detected from oligodendrocytes is low compared to prior time-points when acquired at the same settings. As a result, the BOSS model reported marked loss of cells at T=7 (fourth column) and a drop in the number of tracked oligodendrocytes at T=8 (first column) as shown in Figure S4. In this regard, control images can be used to test quality of fluorescence detection for each individual time-point, and allow rejection of images with poor signal-to-noise for analysis. Our BOSS model can also be used to detect any irregularities that arise over longitudinal experiments - comparison to prior images may prompt immediate assessment of image acquisition parameters in real time. Alternately, BOSS model code can potentially be modified to correct for differences in signal-to-noise ratio at individual time-points.

The BOSS model assumes that the images have been processed to be spatially registered longitudinally, as the CCL annotates the oligodendrocytes based on the location. If images are misaligned, which may occur due to 3-dimensional movement of cells within the brain itself^6^ then the CCL fails to recognize the oligodendrocytes that exist over time and instead reports false lost oligodendrocytes and new oligodendrocytes. Due to misalignment in some cases, BOSS reported a similar number of tracked oligodendrocytes to blinded human observers but reported higher numbers of new/lost oligodendrocytes. In the future, extensions of the BOSS model to incorporate automated longitudinal alignment in the tracking algorithm could aid the BOSS model to align the images accordingly.

Cell segmentation and quantification is a crucial part of the analysis of two-photon microscopy data, but was a significant barrier because manual curation by blinded human observers is costly, time-consuming, and not scalable to large datasets. Here, we proposed a model that allows automated quantification that can give robust, reliable, and reproducible results. We expect our BOSS model to be able to solve an important bottleneck in cell imaging research as our model does not require fully curated data (training data), reduces computational time, and most importantly recapitulates cell dynamic patterns as measured a blinded human observer. Our work is a significant step forward in automated segmentation and tracking of ROI in longitudinal three-dimensional microscopy images, and we expect the BOSS model to facilitate research in this area.

## 4 Methods

We developed an unsupervised segmentation system for annotating and registering oligodendrocytes in three-dimensional images (location denoted by (*i, j, k*), where *i* = 1, …, *I, j* = 1, …, *J, k* = 1, …, *K*) across multiple time-points (time-point denoted by *t*, where *t* = 1, …, *T*). We annotated cells using three main stages: (1) intensity modeling by fitting a mixture of beta distributions; (2) noise filtering by a median filter; and (3) annotation and registration by connected component labeling. The pipeline of the BOSS model is presented in Figure 5.

**Figure 5.**
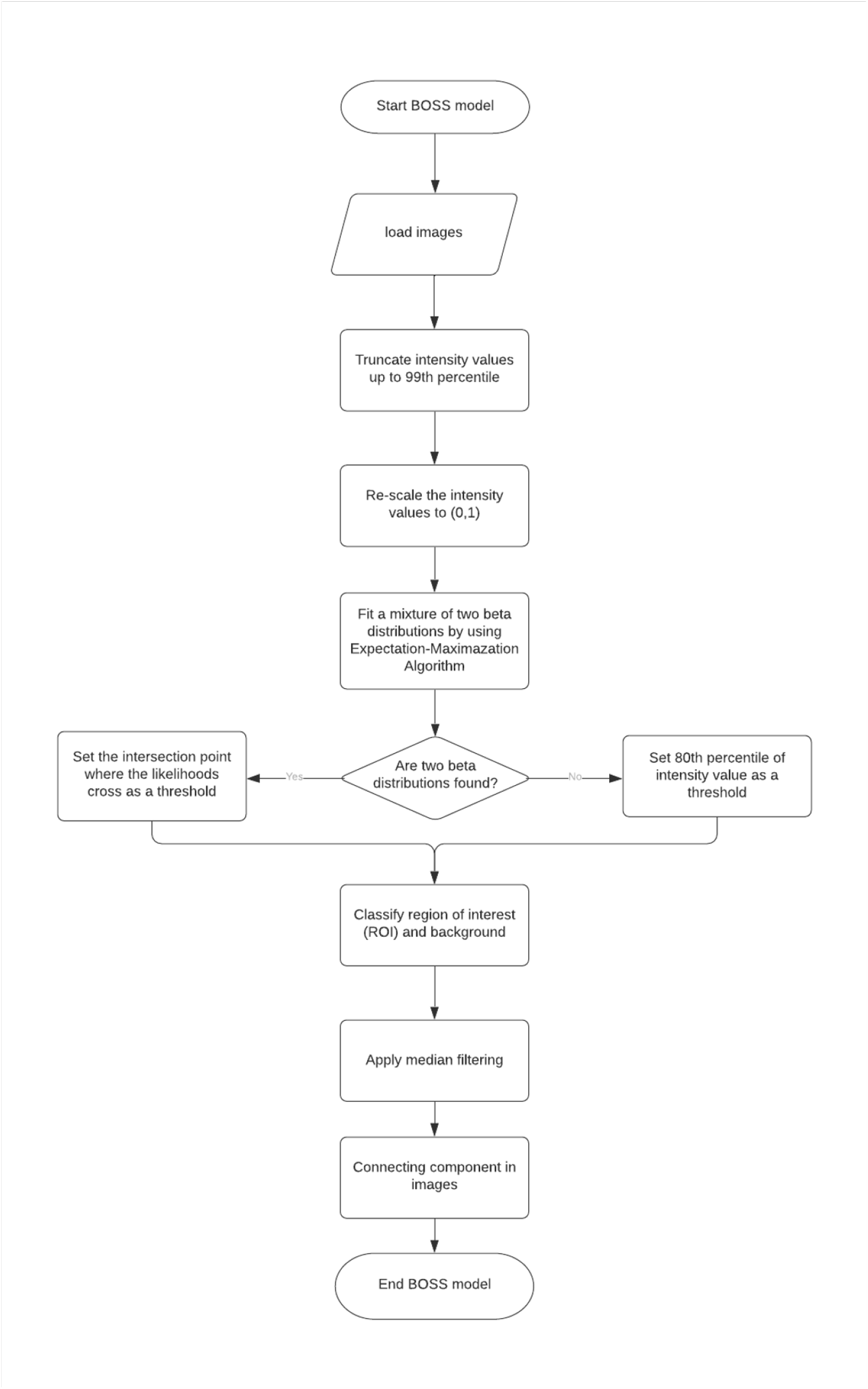
Pipeline of the BOSS model. The BOSS model takes longitudinally aligned images as input. With pre-determined parameters, the BOSS model labels voxels under a threshold as background. After re-scaling the intensity value to [0,1], a mixture of two beta distributions is fit. If two beta distributions are fit, then set the intersection point as a threshold to label the region of interest (ROI) and background. If only one beta mixture component is detected, then voxels under a specified threshold are labeled as background, and those above the threshold are labeled as cell bodies. Then, a median filter is applied to denoise the ROI and oligodendrocytes are labeled using connecting component labeling (CCL).

### 4.1 Image Acquisition and Preprocessing

Throughout this article, we used eight longitudinal image stacks that had been registered using FIJI plugin ‘Correct 3D Drift’^15^, from *n*=3 cuprizone-treated mice and *n*=2 control mice^6^. There were two regions each from two of the cuprizone-treated mice, one region from one cuprizone-treated mouse, two regions from one control mouse and one region from the other control mouse. The size of the images were 425 µm x 425 µm x 550 µm and measured in 1024 × 1024 × 190 voxels for analysis. Images were obtained every 1-2 weeks and followed through 10 weeks (9 time-points) for cuprizone mice and 9 weeks (8 time-points) for control mice.

### 4.2 Intensity Modeling

*In vivo* two photon fluorescence microscopy allows visualization of oligodendrocytes with high resolution. However, the resolution and brightness deteriorates in deeper layers. To account for heterogeneous intensity across cortex depth, we parameterized the intensity of each slice as a two-dimensional image.

The observed intensity of the original images was bounded in [0,255]. We first re-scaled the intensities to [0,1]. Then we truncated up all pixels below the 99th percentile as most of the pixels were part of the image background. As the support of the intensity is bounded, we binarized the two-dimensional images by fitting a mixture of two beta distributions.

Let Y_*i, j,k,t*_ indicate the random variable of intensity value located at {*i, j, k*} and time-point *t* and **Y**_*i, j*_ indicate the collection of image intensity values in two-dimensional (*i, j*) image for fixed *k* and *t* with probability density function *f*.

We modeled Y_*i, j,k,t*_ (hereafter, abbreviated as Y) as coming from one of two different beta distributions *f*_0_, *f*_1_ with proportions *p*_0_ and *p*_1_. Let Z_*i, j,k,t*_ (hereafter, abbreviated as Z) indicate the latent variables representing the mixture component of Y such that P(Z = 0) = *p*_0_ and P(Z = 1) = *p*_1_ = 1 − *p*_0_ where Z∈{ 0, 1} at {*i, j, k}* and time-point *t* and **Z**_*i, j*_ indicate the collection of latent variables in two-dimensional (*i, j*) image for fixed *k* and *t*. Define a random variable Y conditional on the latent variable Z such that Y ∼*f*_0_ if Z = 0 and Y∼ *f*_1_ if Z = 1. Then, the conditional density *f* (Y| Z = *z*) = *f*_*z*_ is the beta distribution for the *z*-th class. The joint probability density function of Z and Y is then given by *f* (Z = *z*, Y = *y*) = *p*_*z*_ *f*_*z*_(*y*). The marginal density of **Y**_*i, j*_ is given as

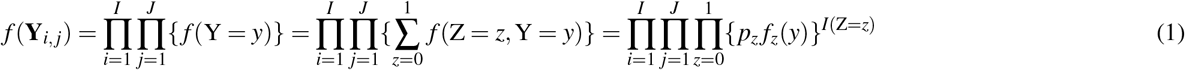

where

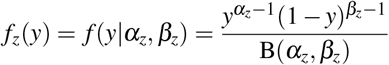

for *z* = 0,1 and 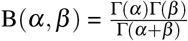, where Γ(·) is the gamma function. Our goal is to estimate unknown parameters, *α*_0_, *α*_1_, *β*_0_, *β*_1_ and *p*_0_, to maximize the *f* (**Y**). We estimated these unknown parameters by implementing an expectation maximization (EM) algorithm.^11^

Let *θ* = (*α*_0_, *α*_1_, *β*_0_, *β*_1_, *p*_0_) indicate the vector of unknown parameters that we want to estimate and *θ* ^(*r*)^ indicate the current value of parameters, *θ* ^(*r*+1)^ indicate the updated parameters. Let *L*(*θ* | **Y**_*i, j*_, **Z**_*i, j*_) and *l*(*θ* | **Y**_*i, j*_, **Z**_*i, j*_) indicate the likelihood and log-likelihood function for complete data. We can formulate the likelihood as

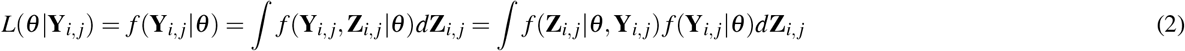

As we do not observe the **Z**_*i, j*_, we implemented an EM algorithm to find the maximum likelihood estimator 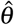 by iterating E- and M-steps as follows:

#### 4.2.1 E-step

Define *Q*(*θ* |*θ* ^(*r*)^) as the expected value of the log-likelihood of *θ* with respect to the conditional distribution of **Z**_*i, j*_ given **Y**_*i, j*_ and current parameters *θ* ^(*r*)^.

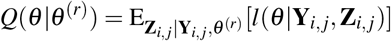

In E-step, we calculate *Q*(*θ* |*θ* ^(*r*)^) as follows.

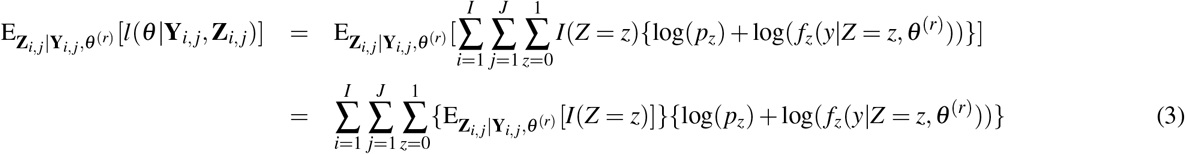

Note that 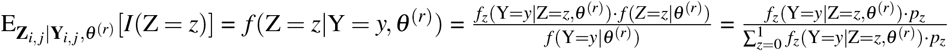

#### 4.2.2 M-step

In the M-step, we update the parameters *θ* ^(*r*+1)^ by maximizing *Q*(*θ* |*θ* ^(*r*)^) such that

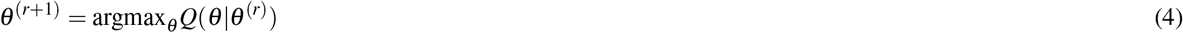

There is no closed form solution to maximize *Q*(*θ θ* ^(*r*)^), so we used a numerical approximation function, *stats::optim()*, in R to find the maximum likelihood estimators.

For initialization, we randomly assigned *Y* to belong either *Z* = 0 or *Z* = 1, as has been found to be efficient^22^. If the EM algorithm converges with two beta distributions, then we retrieved the intersection point where the likelihoods cross and used this for thresholding. If the EM algorithm converges with one beta distribution, then we retrieved the 80th percentile of pixel intensities and used this for thresholding. Note that this 80 percentile value is flexible and the user can determine the optimal percentile based on image quality. Values above the threshold were labeled as region of interest (ROI) and values below this threshold were labeled as background.

Fitting a mixture model took less than 20 seconds per two-dimensional image with dimension [1024 × 1024]. In our analysis, each data set has a dimension of [1024 × 1024 × 190 × 9] or [1024 × 1024 × 190 × 10] and took around 10 hours on a single core run serially. However, this step can be parallelized and computing time can be significantly reduced^23^.

### 4.3 Noise Filtering

Oligodendrocytes have complex processes that extend from the cell body to interact with axons and form myelin sheaths; however, these processes are ancillary to cell body counting. Myelin sheaths in microscopy image is shown as impulsive noise, which has small bright voxels in dark regions. Compared to linear filters (e.g. gaussian filter and mean filter), median filters do not produce image blurring which is desired because image is already binarized to zero and one^24^. We stacked the two-dimensional images that were binarized in Section 4.2 and applied the median filter.^25^

Let 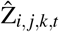 indicate the fitted latent variable of Z_*i, j,k,t*_ by intensity modeling in section 4.2 and let 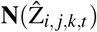indicate the collection of neighboring intensity values of 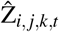. The median filter procedure calculates the median value of the vector 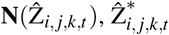

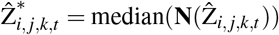

Depending on the size of the cell body of oligodendrocytes, window size of median filter can be optimized. We used the window size of [11,11,3] for fixed *t* as myelin sheaths are less likely to present across depths of the cortex while the cell body are more isotropic. Note that this size is flexible and user can choose the size based on the size of the oligodendrocytes and acquired image resolution.

For computation, median filtering took less than 5 minutes with dimension [1024 × 1024 × 190] in single core and may take 50 minutes to filter full longitudinal three-dimensional image with dimension [1024 × 1024 × 190 × 10]. However, this step again could benefit from parallelization.

### 4.4 Cell Annotation, Registration, and Postprocessing

In graph theory, a connected component is defined as a subgraph in which every two nodes are connected to each other by edges, but not connected to additional nodes in the supergraph. Let graph *G*_*i*_ = {*V*_*i*_, *E*_*i*_} indicate the *i*-th four-dimensional graph with the node set *V*_*i*_ and edge set *E*_*i*_. From binarized, median-filtered intensity value 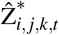located at {*i, j, k*} and time-point *t*, voxel {*i, j, k, t*} is considered node where 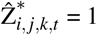while the tracts that connect the nodes of neighboring nodes are considered as edges.

To define the edges in four-dimensional images, we defined the neighborhood as the kernel size [3,3,3,3] such that only elements next to the 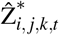 is considered as neighborhood. The resulting neighboring value, 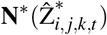, is 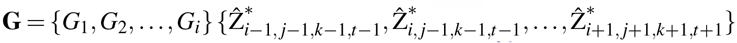. Then, we determined the number of connected components **G** = {*G*_1_, *G*_2_, …, *G*_*i*_} in four-dimensional images ^26^. For images of lesser quality, larger kernel size can be used to accommodate the less spatially

We expect to have a high signal-to-noise image throughout the experiment so that connecting component algorithm can form a subgraph over timepoints. However, if an interim image has low signal-to-noise (minimal detection of emitted signal), subgraphs are divided and therefore the BOSS model reports two cells, one lost cell at the time-point of the low signal-to-noise image time-point and one new cell at the following time-point. To adjust the heterogeneous resolution of the interim image, we assumed that the subgraph located at the neighborhood of {*i, j, k*} over time-point *t* to be considered as the same subgraph. This kernel of the neighborhood can be optimized. Once **G** is determined, we filtered out the connected components in **G** that have the node sets with less than 30 nodes. This enabled the algorithm to keep the cell body while reducing noise. This number is also a tuning parameter that is flexible, and larger or small size can be used based on the relative size of oligodendrocytes. Since the operation is sequential on four-dimensional array, parallelization is not available in this step.

---

**Figure S2.**
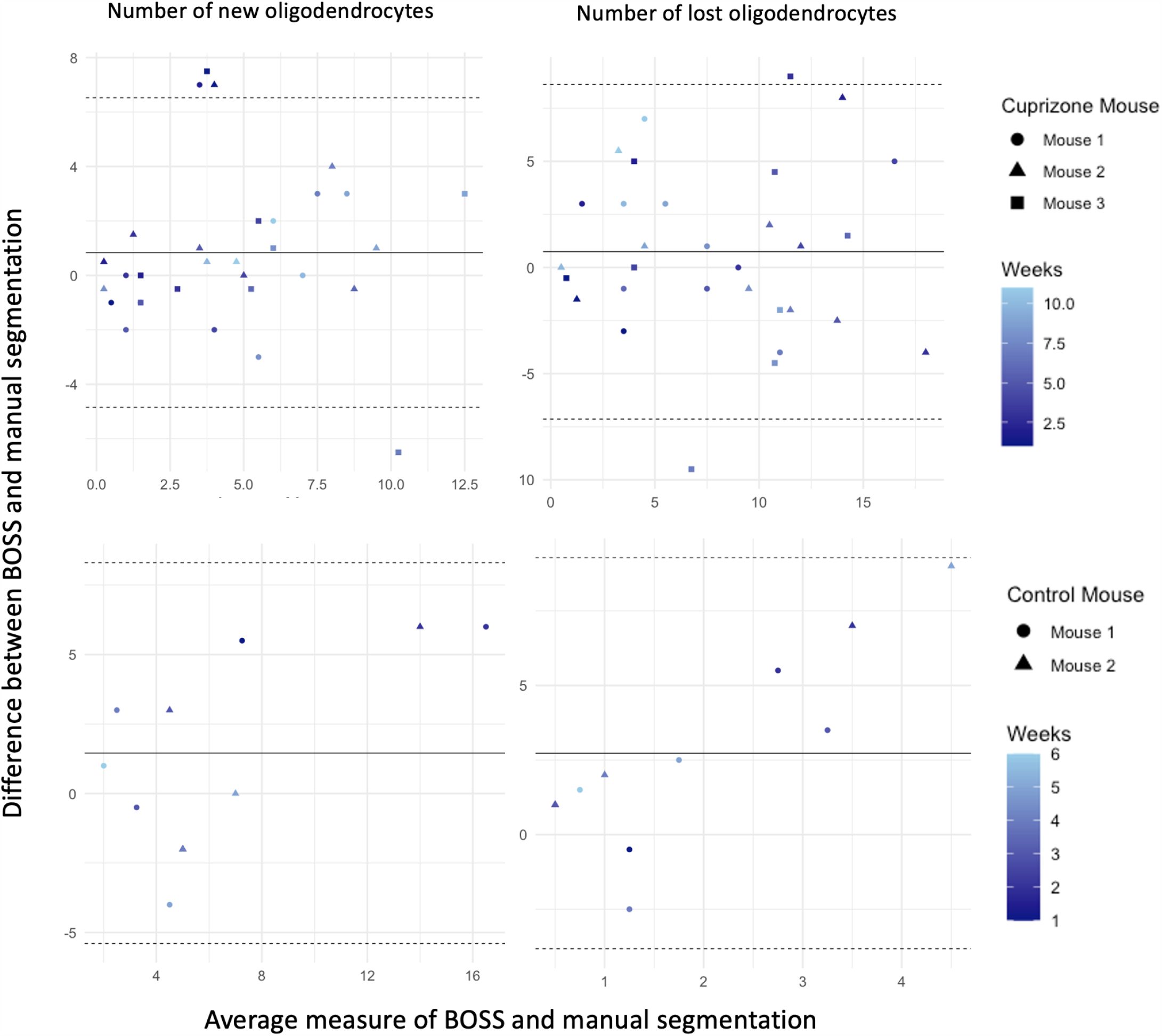
Bland–Altman (BA) plot of the new oligodendrocytes (first column) and lost oligodendrocytes (second column). Y-axis indicates the difference between BOSS and manual segmentation and X-axis indicates the average measure of BOSS and manual segmentation. Straight bold line in BA plot indicates the mean difference and dashed line indicates the mean difference plus and minus 1.96 times the standard deviation of the differences.

**Figure S3.**
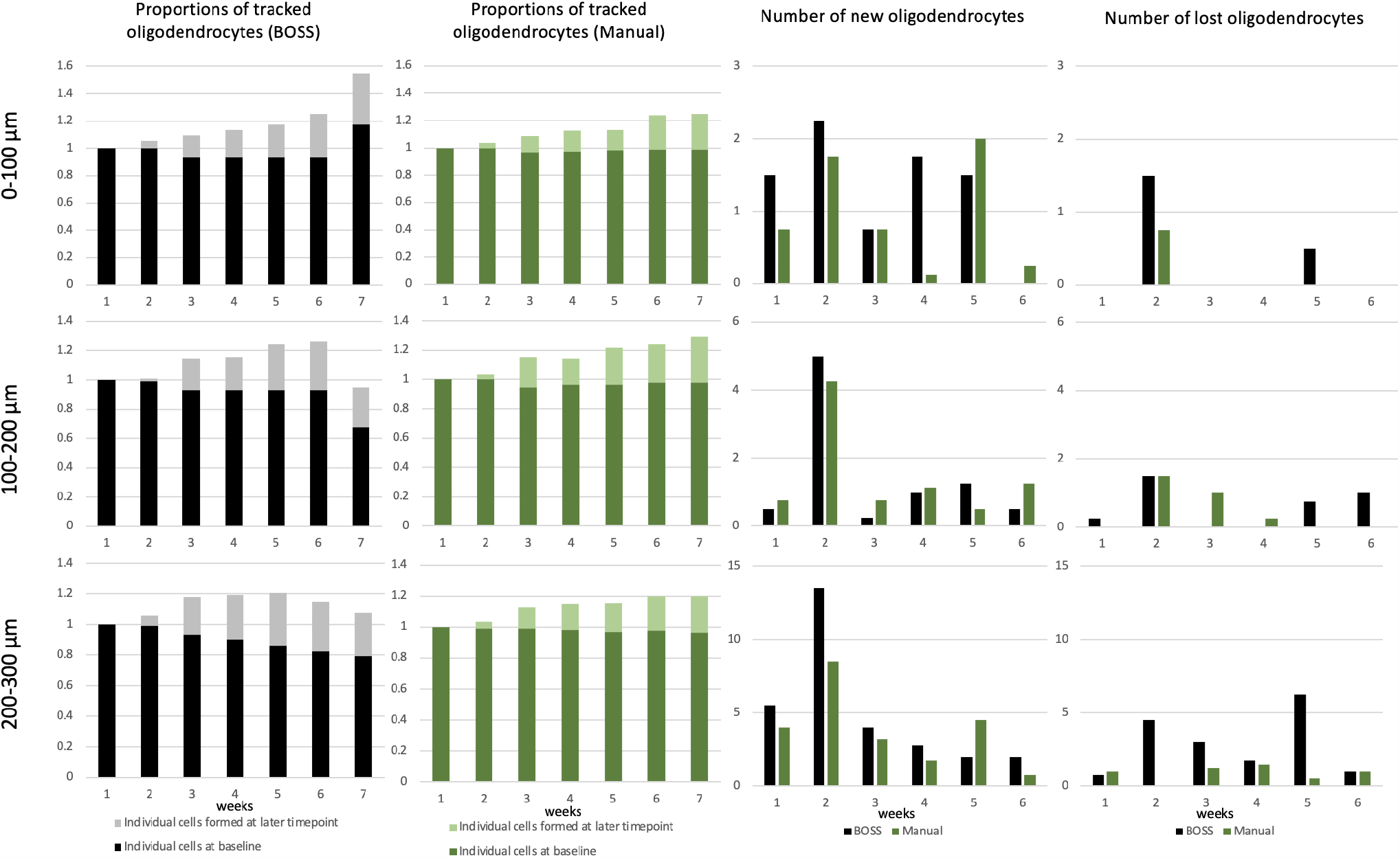
Result of the BOSS model across time points, stratified by depth in somatosensory cortex for control mice. The rows indicate the 0-100 µm, 100-200 µm, and 200-300 µm depth regions of the images. The first column shows the average number of tracked oligodendrocytes by the BOSS model. The first column shows the average number of tracked oligodendrocytes by the BOSS model. The second column shows the average number of tracked oligodendrocytes by blinded human observer. The third column shows the average number of newly formed oligodendrocytes. The fourth column shows the average number of lost oligodendrocytes.

**Figure S4.**
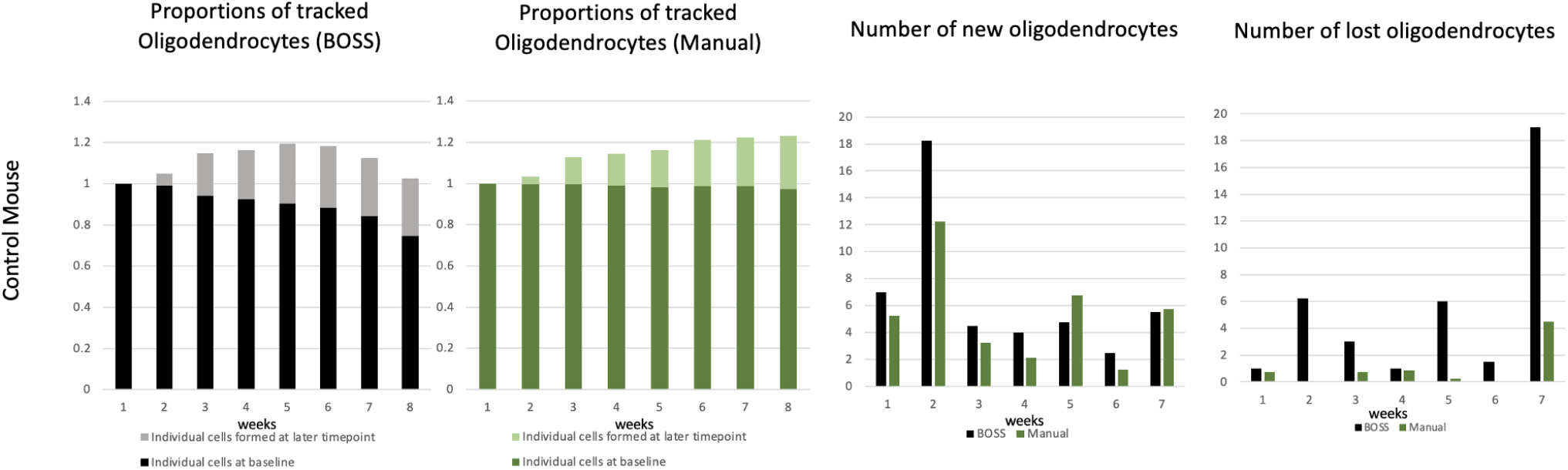
Results of the BOSS model across time-points when applied to the control mice including under-exposed eighth time-point. The first/second column shows the proportions of the number of tracked oligodendrocytes by the BOSS model/blinded human observer. The third column shows the average number of newly formed oligodendrocytes at each time-point. Each time-point represents the number of new oligodendrocytes observed at its next time-point. The fourth column shows the average number of lost oligodendrocytes at each time-point. Each time-point represents the number of lost oligodendrocytes observed at its next time-point. The BOSS model reports notable loss of oligodendrocytes at seventh time-point.

**Figure S5.**
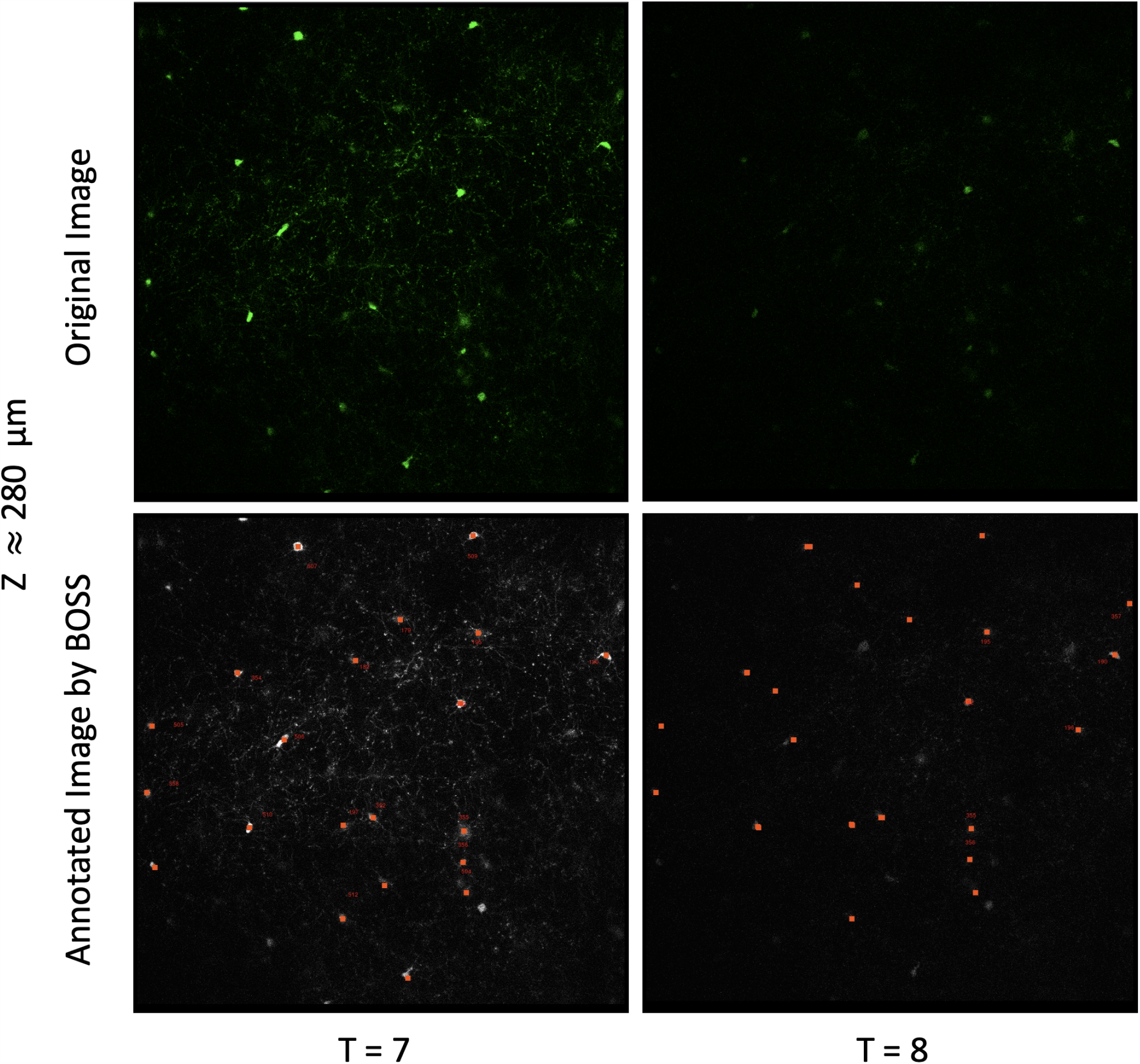
Original images in the first row and annotated images by the BOSS model in the second row. First and second columns show the original/annotated image at T=7 (normal) and 8 (under-exposed). The index number next to the red rectangular is the unique identifier that given by the BOSS model. Red rectangles with index numbers in the annotated image indicate that oligodendrocytes are annotated at both the seventh and eighth time points. Red rectangles without index numbers indicate that oligodendrocytes are annotated at the seventh time-point but lost at eighth timepoint by the BOSS model. The original image at T=8 has low signal-to-noise and the BOSS model did not annotate the oligodendrocytes (index numbers exist at T=7 but not at T=8) and therefore reports many lost oligodendrocytes.

